# Dissection of genotype-by-environment interaction and simultaneous selection for grain yield and stability in faba bean (*Vicia faba* L.)

**DOI:** 10.1101/2022.09.08.507215

**Authors:** Tadesse S. Gela, Hamid Khazaei, Rajib Podder, Albert Vandenberg

## Abstract

Increasing faba bean production is indispensable to supply the growing demand for plant-based protein on the global scale. A thorough understanding of genotype (G) × environment (E) interaction (GEI) patterns is critical to developing high-yielding varieties with wider adaptation. Thirteen faba bean genotypes were evaluated in 15 environments during 2019–2020 in western Canada to estimate their yield stability using different stability statistics. The combined analysis of variance and additive main effects and multiplicative interaction (AMMI) analysis revealed that G, E, and GEI effects were highly significant (*P*<0.001), indicating differential responses of the genotypes across the environments, enabling the stability analysis. The result of the model comparison found the best linear unbiased prediction (BLUP) to outperform AMMI models. The BLUP-based biplot of the weighted average of absolute scores (WAASB) stability and mean grain yield identified AO1155 (Navi), 1089-1-2, 1310-5, DL Tesoro, and 1239-1 as high-yielding and stable genotypes. The correlation analysis revealed that most of the stability parameters had a strong association with grain yield and with each other, indicating that they should be used in combination with one another to select genotypes with high yield. Overall, the WAASB superiority index (WAASBY) and the average sum of ranks of all stability statistics identified the same genotypes in terms of high yielding and stability, and genotype AO1155 is considered the most stable and highest yielding among the tested genotypes. Genotypes with stable yields across environments would be beneficial for faba bean genetic improvement programs globally.

**Core Ideas:**

- Stability analysis was estimated using 13 faba bean genotypes over 15 site-years.
- The different stability methods described genotypic performance in different ways.
- The majority of stability models showed a strong rank correlation with grain yield.
- AMMI and BLUP analyses revealed a highly significant G×E interaction, with BLUP outperforming AMMI.
- Overall, the employed stability statistics identified AO1155 as the highest yielding and most stable genotype.

## 1 INTRODUCTION

Faba bean (*Vicia faba* L.) is an important cool-season grain legume crop cultivated worldwide for its high seed protein content and excellent biological nitrogen fixing ability. Its seed protein content is about 30% of the seed dry matter (Griffiths and Lawes, 1978; Khazaei and Vandenberg, 2020), which is highly valuable for human consumption and animal feed (Stoddard et al., 2009; Crépon et al., 2010). It has great potential to contribute to fulfilling the increasing global demand for plant-based protein. The crop integrates well into sustainable agricultural systems due to its capacity to improve soil nitrogen fertility and break the cycle of biotic stress in cereal-based cropping systems (Köpke and Nemecek, 2010; Duc et al., 2015), and extend crop rotations that include other pulse crops that are susceptible to *Aphanomyces* root rot (Moussart et al., 2008). Faba bean can thrive in a wide range of soil types and climates, including cool, moist, and warm temperate and subtropical regions (Duc et al., 2015). Nevertheless, it is known for its unstable yield performance (e.g., Cernay et al., 2015; Reckling et al., 2018) due to its susceptibility to interannual fluctuations and environmental variations (Zong et al., 2019). The main goal of breeding programs is to increase and maintain the productivity of the crop by developing high-yielding and stable varieties. For this reason, breeders test large numbers of genotypes in various environments to evaluate the yield stability and wide adaptability of the genotypes.

Multi-environment trials (METs) have an important role in interpreting the genotype × environment interaction (GEI) effect and selecting superior genotypes at the end of the variety development pipeline. Modeling the GEI in METs can help in determining the adaptability and stability of genotypes across a wide range of environments. However, this concept has been defined in different ways in interpreting GEI (Gauch and Zobel, 1996), resulting in an increasing number of stability statistics (Pour-Aboughadareh et al., 2022a). Huehn (1996) classified stability statistics into parametric and non-parametric methods. The parametric methods, which include univariate and multivariate stability analysis, mainly rely on distributional assumptions about the environment, genotype, and their interaction effects. The non-parametric approaches are estimated based on the mean values of the response variable and ranking of genotypes without any primary distribution assumptions. Accordingly, a genotype is considered stable if its ranking remains largely constant across environments (Flores et al., 1998) and the addition or deletion of one or a few genotypes has no significant effect on the results (Huehn, 1990). Several parametric and non-parametric methods and models have been developed to analyze the extent of GEI and determine the yield performance and stability of genotypes (reviewed by Pour-Aboughadareh et al., 2022a). Each of these methods has its own set of strengths and weaknesses in describing the phenomenon of GEI, so a combination of statistics from both approaches may provide a better understanding of GEI (Becker and Leon, 1988). In faba bean, a similar strategy has been applied (e.g., Temesgen et al., 2015; Sheikh et al., 2021).

The additive main effects and multiplicative interaction (AMMI model; Gauch, 1992) is a multivariate parametric approach that is widely used to analyse and interpret GEI in METs. The AMMI is a hybrid model that employs the analysis of variance for additive or main effects and principal component analysis (PCA) for the multiplicative effects to understand the patterns of GEI (Zobel et al., 1988). The graphical biplot tools of the AMMI model provide an easy interpretation of yield performance and stability simultaneously, as well as the identification of mega-environment (Zobel et al., 1988; Gauch, 1992; Gauch et al., 2008). Previous studies have demonstrated the usefulness of the AMMI model in identifying superior faba bean genotypes in terms of yield performance and stability, as well as its advantage in describing the GEI effect (e.g., Flores et al., 1996; Flores et al., 1998; Fikere et al., 2008; Tadesse et al., 2017). However, the AMMI model has a weakness in analysing the structure of the linear mixed-effect model (LMM). Alternatively, the best linear unbiased prediction (BLUP; Smith et al., 2005) methods were proposed to estimate GEI in METs based on the LMM, which is efficient in the estimation of random effects. In some cases, the BLUP was found to be more reliable than the AMMI model at making predictions (e.g., Piepho, 1994; Olivoto et al., 2019). Consequently, Olivoto et al. (2019) integrated the graphical tools of the AMMI model into the BLUP technique and developed a new stability statistic, WAASB (weighted average of absolute scores). The WAASB has been used to identify high-yielding stable genotypes in many crops (Koundinya et al., 2021; Nataraj et al., 2021; Pour-Aboughadareh et al., 2022b; Yue et al., 2022), but not in faba bean.

Several faba bean cultivars have been developed in western Canada since the late 1970s (Khazaei et al., 2021). However, yield stability has not been studied on modern faba bean cultivars. Overall, to describe the extent of GEI and select high-yielding stable genotypes, a number of approaches, including those listed above, have been proposed. In the current study, a total of 20 stability statistics, including the newly introduced WAASB, were used to investigate the grain yield performance and stability of 13 faba bean genotypes in METs, and the relationships between stability parameters were analyzed. Finally, the average sum rank of all the stability statistics was computed and compared with the WAASB superiority index (WAASBY) for simultaneous selection of grain yield performance and stability of faba bean genotypes.

## 2 MATERIALS AND METHODS

### 2.1 Plant material and growing conditions

A set of thirteen white-flowered (low tannin) faba bean genotypes, including ten advanced breeding lines and three check varieties (cv. Snowbird, DL Tesoro, and DL Rico), were grown at 12 locations in western Canada during the 2019 and 2020 cropping seasons. The test genotypes originated from different major faba bean breeding programs in Europe and Canada (Table 1). The locations were representative of typical faba bean growing regions in the dark gray, black, and dark brown soil-climatic zones of western Canada. Each year and location was treated as a separate environment, making a total of 15 environments: seven in Manitoba, six in Saskatchewan, and two in Alberta, Canada. The trials were sown in May and harvested from September to November. Climatic information was retrieved from the Environment Canada database for the weather station nearest to the field site. Detailed characteristics of the test locations are presented in Table 2.

**Table 1.** Information on the tested faba bean genotypes tested in this study

| Genotype code | Name | Vicin-convicine | Breeder \ Origin |
| --- | --- | --- | --- |
| G1 | Snowbird | High | Limagrain Advanta, The Netherlands |
| G2 | DL Tesoro | High | NPZ, Germany |
| G3 | DL Rico | Low | NPZ, Germany |
| G4 | NPZ 16.7610 | Low | NPZ, Germany |
| G5 | NPZ 16.7601 | Low | NPZ, Germany |
| G6 | AO1155* | Low | Agri Obtentions, France |
| G7 | 951-1-11 | Low | CDC, Usask, Canada |
| G8 | DL 18.7602 | Low | NPZ, Germany |
| G9 | DL 18.7603 | Low | NPZ, Germany |
| G10 | DL 18.7604 | Low | NPZ, Germany |
| G11 | 1089-1-2 | Low | CDC, Usask, Canada |
| G12 | 1310-5 | Low | CDC, Usask, Canada |
| G13 | 1239-1 | Low | CDC, Usask, Canada |

CDC, Crop Development Center; Usask, University of Saskatchewan; NPZ, Norddeutsche Pflanzenzücht, Germany. AO1155 is registered as “Navi” in western Canada (https://inspection.canada.ca/english/plaveg/pbrpov/cropreport/faba/app00012063e.shtml)

**Table 2.**
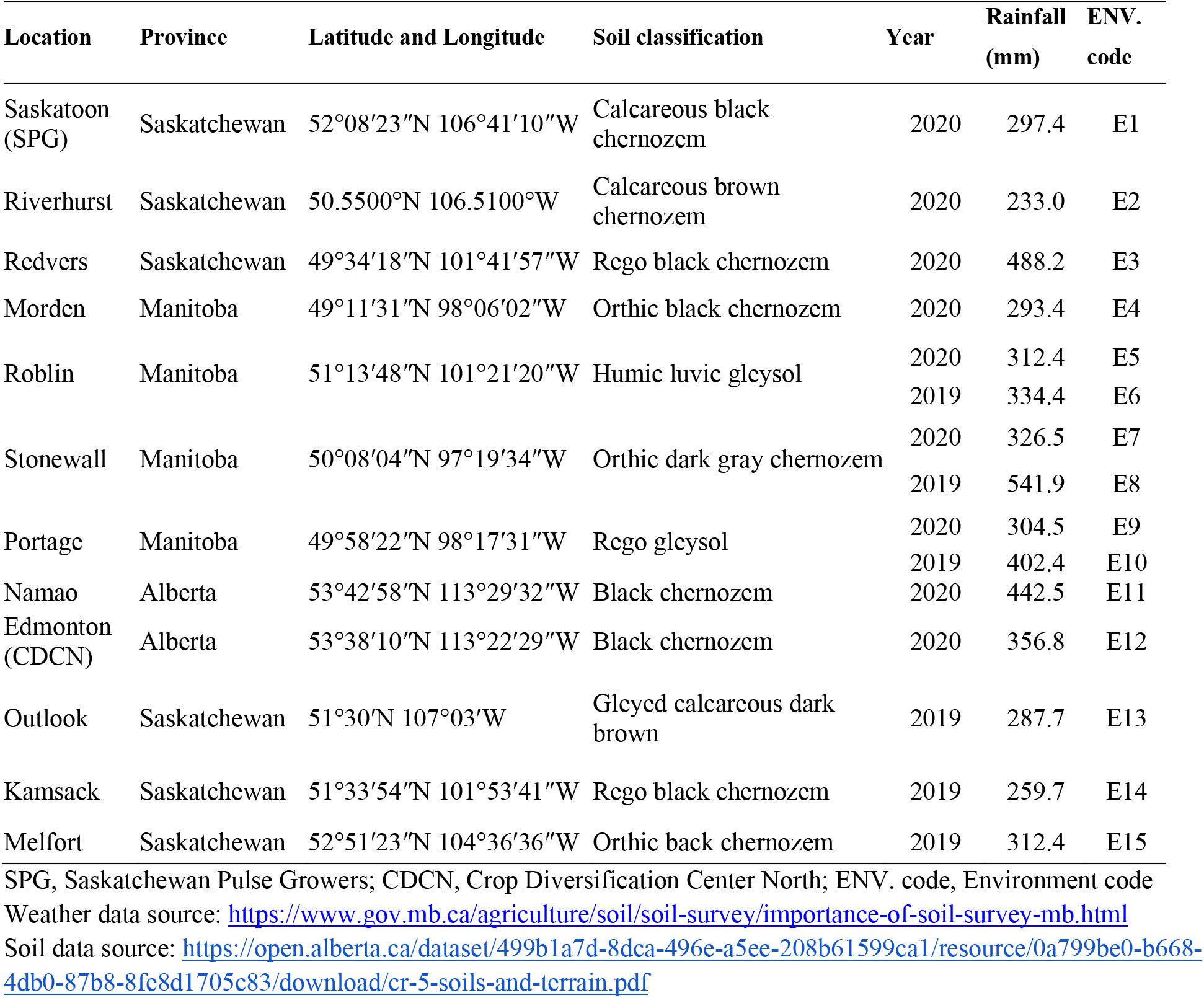
Field plot locations, soil classification and rainfall over the growing season 2019-2020

### 2.2. Experimental design

The field experiments were conducted in a randomized complete block design (RCBD) with three replications, each in a 1.2 m × 5 m plot. All necessary crop management practices such as nutrients, weed control, and pesticide applications were followed as per recommended practices at each location. These agronomic practices were applied uniformly to the entire experimental area and were treated as non-experimental variables. Plots were harvested with combine harvesters and seeds were cleaned to determine grain yield and seed weight at 10% standard grain moisture content for data analysis.

### 2.3. Statistical analyses

The analysis of variance (ANOVA) of grain yield data from each environment (combination of years and locations) was first analysed separately. The combined ANOVA was then performed to determine the effects of genotype, environment, and genotype by environment interaction. Furthermore, the AMMI analysis was conducted to partition GEIs into different principal components. All these statistical analyses were performed in R software 4.0.3 (R Core Team, 2020) using the package “metan” (Olivoto and Lucio, 2020). The combined and AMMI ANOVA was performed using a linear model with an interaction effect, whereas the variance components were estimated in a linear mixed-effect model using restricted maximum likelihood (REML) considering genotype and genotype-vs-environment as random effects.

The AMMI family model and the BLUP model were tested for prediction accuracy by comparing their root mean square prediction difference (RMSPD) estimates (Piepho, 1994). Likewise, the WAASB (the weighted average of absolute scores from the singular value decomposition of the matrix of best linear unbiased predictions for the GEI effects generated by a linear mixed-effect model) statistics was employed to analyse the stability (Olivoto et al., 2019). This method integrates AMMI and BLUP model features for identifying highly productive and stable genotypes. Correspondingly, the superiority index of WAASB was calculated for simultaneous selection for yield and stability by weighting the WAASB stability value and mean yield performance (Y) (WAASBY, Olivoto et al., 2019).

Additionally, eleven parametric and nine non-parametric common stability statistics were calculated (see Table 3), and furthermore, the investigated genotypes were ranked based on each statistic. All these stability statistics were estimated using a web-based STABILITYSOFT program (Pour-Aboughadareh et al., 2019) and the “metan” package in R. An overview of their equations and how they were used in the analysis of GEI effects is given in a recent review by Pour-Aboughadareh et al. (2022a). Spearman’s rank correlation was computed to detect the association between the calculated stability statistics using the “corrplot” package in R. To better understand the interrelationships among stability measures, PCA was performed using the “factoextra” package in R. Moreover, to group the investigated faba bean genotypes into similar mean yield and stability clusters, a hierarchical cluster analysis (HCA) was computed based on average sum of ranks (ASR) of all stability measures and mean yield through Ward’s method and Euclidean distance as a dissimilarity measure using the “ggdendro” packages in R.

**Table 3.**
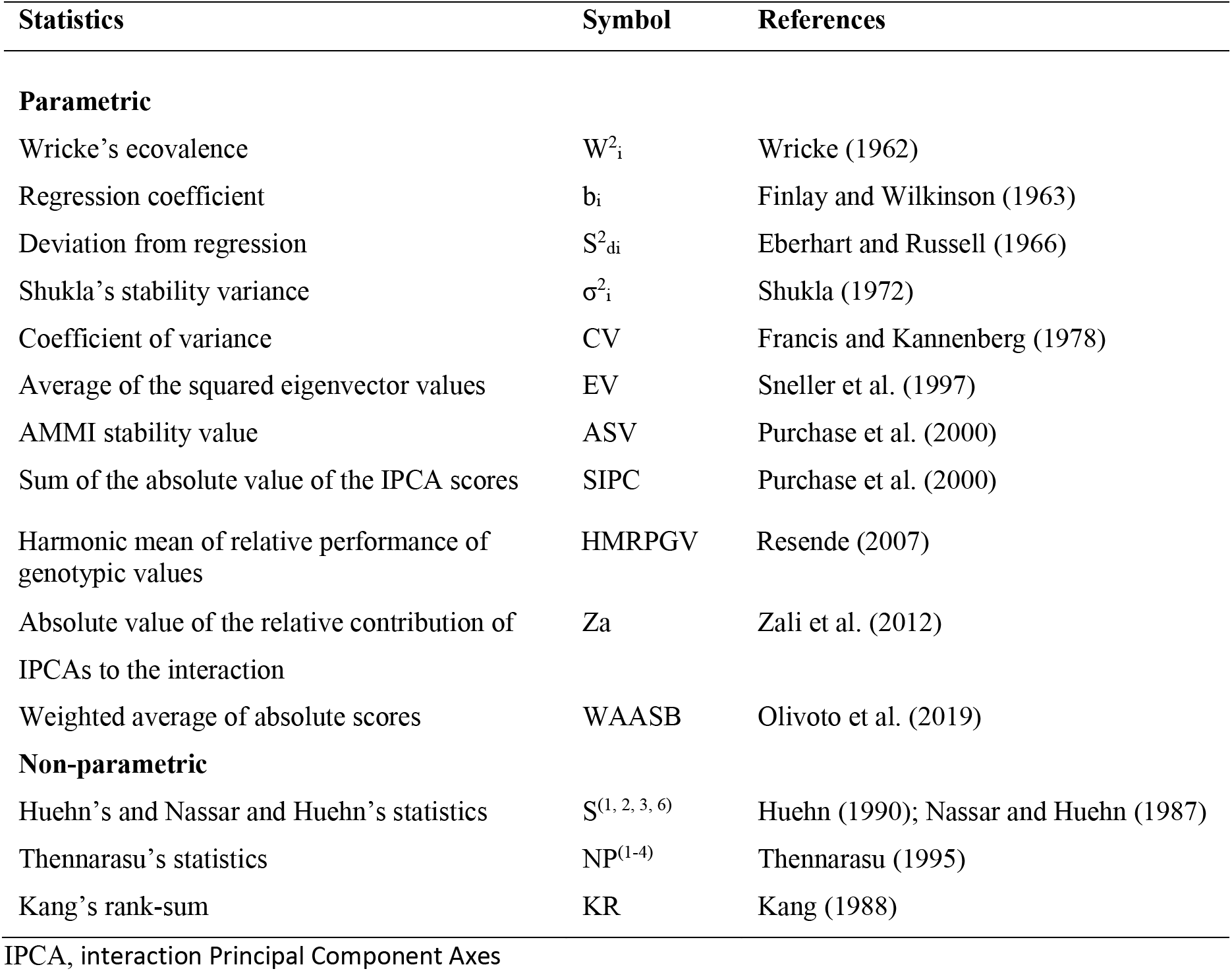
List of parametric and non-parametric stability statistics computed in this study

## 3 RESULTS

### 3.1 Analysis of variance and mean performances

The analysis of variance of grain yield for the individual environments revealed significant differences for genotype effects (*P*<0.05), except for E5 (Roblin 2020) and E13 (Outlook 2019). The value for coefficient of variation ranged from 5.46 to 15.31 (Supplemental Table S1), indicating the suitability of the data for further analysis. The joint ANOVA across the 15 test environments showed the presence of highly significant variation (*P*<0.001) for the main effects due to genotype, environment, and their interaction (Table 4). The analysis reported that the highest proportion of variance was contributed by environment effect (76.6%), followed by genotype by environment interaction effect (9.5%) and genotype effect (4.2%). The mean grain yields of environments varied from 0.881 t ha^-1^ (E10, Pontage 2019) to 3.514 t ha^-1^ (E13, Outlook 2019), with a grand mean of 2.301 t ha^-1^. The highest mean grain yield (2.534 t ha^-1^) was produced by G11 (1089-1-2) and the lowest (2.030 ha^-1^) was produced by G9 (DL 18.7603) with a grand mean of 2.301 t ha^−1^ (Supplemental Table S2).

**Table 4.** Analysis of variance for combined and AMMI analysis for grain yield of 13 faba bean genotypes evaluated at 15 environments during 2019–2020 cropping seasons

| Sources of variation | DF | Sum square | Mean square | TSS (%) | GEI expl. (%) | Cumulative (%) |
| --- | --- | --- | --- | --- | --- | --- |
| **Combined analysis** |  |  |  |  |  |  |
| Environment (E) | 14 | 288.91 | 20.64*** | 76.58 |  |  |
| Replication / E | 30 | 9.64 | 0.32*** | 2.56 |  |  |
| Genotype (G) | 12 | 15.73 | 1.31*** | 4.17 |  |  |
| GE interaction | 168 | 35.75 | 0.21*** | 9.48 |  |  |
| Residuals | 355 | 27.22 | 0.08 |  |  |  |
| CV (%) |  | 12.07 |  |  |  |  |
| **AMMI analysis** |  |  |  |  |  |  |
| Environment (E) | 14 | 288.91 | 20.64*** |  |  |  |
| Replication / E | 30 | 9.64 | 0.32*** |  |  |  |
| Genotype (G) | 12 | 15.73 | 1.31*** |  |  |  |
| GE interaction | 168 | 35.75 | 0.21*** |  |  |  |
| IPC1 | 25 | 9.65 | 0.39*** | 2.33 | 26.40 | 26.4 |
| IPC2 | 23 | 9.03 | 0.39*** | 2.18 | 24.70 | 51.1 |
| IPC3 | 21 | 7.45 | 0.35*** | 1.80 | 20.40 | 71.5 |
| IPC4 | 19 | 3.61 | 0.19*** | 0.87 | 9.90 | 81.3 |
| IPC5 | 17 | 3.12 | 0.18** | 0.75 | 8.50 | 89.9 |
| IPC6 | 15 | 1.31 | 0.09<sup>ns</sup> | 0.32 | 3.60 | 93.5 |
| IPC7 | 13 | 1.20 | 0.09<sup>ns</sup> | 0.29 | 3.30 | 96.7 |
| IPC8 | 11 | 0.44 | 0.04<sup>ns</sup> | 0.11 | 1.20 | 98.0 |
| IPC9 | 9 | 0.33 | 0.04<sup>ns</sup> | 0.08 | 0.90 | 98.9 |
| IPC10 | 7 | 0.24 | 0.03<sup>ns</sup> | 0.06 | 0.70 | 99.5 |
| IPC11 | 5 | 0.16 | 0.03<sup>ns</sup> | 0.04 | 0.40 | 100 |
| IPC12 | 3 | 0.01745 | 0.00582 | 0 | 0.05 | 100 |
| Residuals | 355 | 27.22 | 0.08 |  |  |  |

\*\* and \*\*\*, significant at the 0.01 and 0.001 probability levels, respectively; ns, non-significant; TSS, total sum of squares; GEI expl., genotype × environment interaction explained; CV, coefficient of variation; DF, degrees of freedom

### 3.2 AMMI Analysis of variance

The multivariate stability model, AMMI analysis for grain yield of the 13 faba bean genotypes tested in 15 environments is presented in Table 4. This analysis shows that the faba bean grain yield was significantly (*P*<0.001) affected by changes in environment, followed by G × E interaction and genotypic effects. The variance explained by the GEI effect was two times greater than the genotype effect. This demonstrates that the genotypes responded differently across environments and that additional stability analysis may be required to fully understand the magnitude of the GEI effect. The GEI effect was further partitioned into 12 interaction principal components (IPCs). The first five IPCs were found to be significant (*P*<0.001) and explained 89.9% of the variation affected GEIs, with a proportion of 26.4, 24.7, 20.4, 9.9, and 8.5% for IPC1-5, respectively (Table 4). The distribution of faba bean genotypes and test environments based on biplots of AMMI1 (mean grain yield × IPC1 scores) and AMMI2 (IPC1 × IPC2 scores) are found in Supplemental Figure S2 A and B. However, as IPC1 and IPC2 were unable to account for 48.9% of the GEI variance, using this information to interpret the GEI could be deceptive. So, to thoroughly analyse the GEI effect, methodologies incorporating more than the first two IPC would be necessary.

### 3.3 Model accuracy testing and predicted means

The grain yield prediction accuracy of the BLUP and AMMI models is tested using RMSPD. The estimated values for each model member family are compared based on the mean of 1000 resampling cycles of RMSPD for each model tested. The model with the smallest RMSPD value is regarded as the most accurate prediction, and vice versa. In the present study, compared to the AMMI family models, the BLUP had the smallest RMSPD followed by AMMI3 (Supplemental Figure S3). Therefore, the BLUP was found to be the most accurate predictive model for faba bean grain yield. The lowest predicted mean was estimated for G9 (2.074 t ha^−1^) and the highest predicted mean was for G11 (2.496 t ha^−1^), with a BLUP mean of 2.301 t ha^−1^ (Figure 1A and Supplemental Table S3). The observed and predicted mean for the genotypes are close to each other, as indicated by the higher value of the genotypic accuracy of selection (As = 0.915, Table 5). Seven genotypes had a mean value greater than the grand mean, and the remaining six genotypes scored below the BLUP mean.

**Figure 1.**
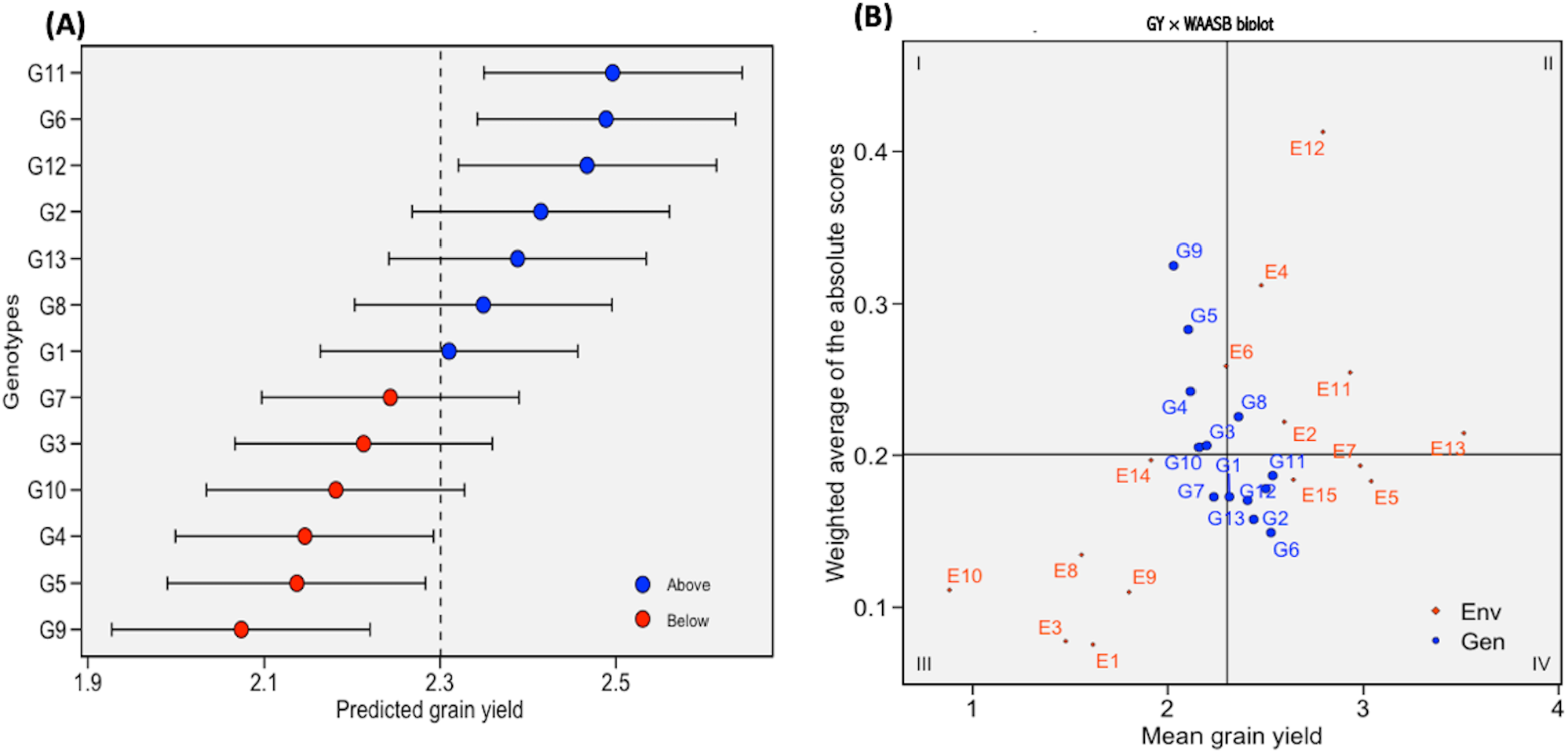
(A) the predicted grain yield performance of 13 faba bean genotypes estimated using BLUP (best linear unbiased prediction). The vertical dotted line indicates the grand mean and the horizontal error bars indicate the 95% confidence interval when considering the two-tailed t-test. (B) biplot of mean grain yield and weighted average of absolute scores for the best linear unbiased predictions of the genotype vs. environment interaction (WAASB). See Tables 1 and 2 for genotypes and environments’ legends

**Table 5.** Estimated variance components and genetic parameters for grain yield of 13 faba bean genotypes evaluated in 15 environments

| Statistics | Likelihood ratio test |  |
| --- | --- | --- |
|  | G | GEI |
| $\chi^2$ | 31.525 | 68.490 |
| *p-value* | 1.97E-08 | 1.28E-16 |
| REML<sup>λ</sup> | Variance components |  |
|  | Estimates |  |
| $\sigma^2_g$ | 0.025 (16.85%)<sup>Ω</sup> | |
| $\sigma^2_{gei}$ | 0.047 (31.74%) | |
| $\sigma^2_e$ | 0.076 (51.42%) | |
| $\sigma^2_p$ | 0.148 | |
| $h^2$ | 0.169 | |
| $R^2_{gei}$ | 0.317 | |
| $h^2_{mg}$ | 0.838 | |
| AS | 0.915 |  |
| $r_{ge}$ | 0.382 | |
| $CV_g$ (%) | 6.877 | |
| $CV_r$ (%) | 12.008 | |
| $CV_g / CV_r$ ratio | 0.573 | |
| SD | 0.810 |  |
| SE | 0.030 |  |

G, genotype; GEI, genotype by environment interaction; <sup>λ</sup>REML, restricted maximum likelihood; $\sigma^2_g$ , genotypic variance; $\sigma^2_{gei}$ , genotype by environment interaction variance; $\sigma^2_e$ , residual variance; $\sigma^2_p$ , phenotypic variance; $h^2$ , broad-sense heritability; $R^2_{gei}$ , coefficient of determination of the interaction effects; $h^2_{mg}$ , heritability of the genotypic mean; AS, accuracy of selection; $r_{ge}$ , genotype –environment correlation; $CV_g$ %, genotypic coefficient of variation; $CV_r$ %, residual coefficient of variation; $CV$ ratio, ratio between genotypic and residual coefficient of variation; SD, standard deviation; SE, standard error. <sup>Ω</sup> Parenthetical values indicate the percentage of the observed phenotypic variance ( $\sigma^2_p$ )

### 3.4 Estimated variance components

Based on the mixed model likelihood ratio test, both genotype and genotype by environment interactions had highly significant (*P*<0.001) effects (Table 5). As illustrated in Supplemental Figure S1B, the GEI was a crossover type (qualitative) and the rank order of the genotypes changed across the environments. Of the phenotypic variance (σ^2^_p_) estimated, the residual variance (σ^2^_ε_) accounted for 51.4%, the GEI (σ^2^_ge_) for 31.7% and genotypic variance (σ^2^_g_) for 16.8% (Table 5). As a result, low broad-sense heritability (h^2^ = 0.169) was observed. The correlation between predicted and observed genotypic values was high (genotypic accuracy of selection, As = 0.915). In contrast, the low correlation between genotypic values across environments (r_ge_ = 0.38) was explained by the high residual coefficient of variation (CV_r_%) and residual variance (σ^2^_ε_) relative to genotypic variance.

### 3.5 Integrating AMMI and BLUP models to understand the GEI

In this study, the WAASB statistics were computed to better characterize ideal genotypes based on both mean grain yield and stability. With this method, the stability of the genotypes can be presented graphically using biplots of the WAASB scores. Figure 1B depicts the grain yield × WAASB biplot with quadrants denoting the four classes of genotypes/environments that simultaneously interpret productivity and stability along the environments. The perpendicular line to the horizontal axis indicates the overall mean (2.301 t ha^−1^) and discriminates the genotypes’ performance above and below the grand mean. The first quadrant represents the most unstable genotypes that have the largest role in GEI and environments with high discrimination ability. Low-yielding genotypes such as G9, G5, and G4 with mean grain yields of less than the overall mean were included. However, no environments were placed in this quadrant. The second quadrant is defined by its highly productive and unstable genotypes along with environments that have good discrimination powers. It included G8, which had a higher grain yield than the overall mean yield, as well as E2, E4, E11, E12, and E13. These environments require special consideration as they discriminate against the high-yielding genotypes. The third quadrant included environments E1, E3, E8, E9, E10, and E14, as well as genotype G7. The genotypes in this quadrant are considered low-yielding and better stable (widely adapted) due to low WAASB scores. These environments can also be regarded as less productive and having a lower ability for genotype differentiation. The fourth part of the biplot comprised G2, G6, G11, G12, and G13, along with E5, E7, and E15. The genotypes within this part have high yield performance, are widely adapted, and the most stable, making them the most desirable genotypes. The environments included in this quadrant can be considered the most productive but with low discrimination abilities. Moreover, genotype G1 (Snowbird) was placed on the frontier of the third and fourth quadrants and showed yield performance equal to the overall mean and higher stability.

### 3.6 Assessment of yield performance stability

#### 3.6.1 Parametric measures of stability

The first criterion for genotype evaluation was mean grain yield. Based on this parameter, genotypes G11, G6, and G12 had the highest, while G4, G5, and G9 had the lowest mean grain yield (Supplemental Table S2). The joint regression model assesses the stability of each genotype based on the bi and *S*^2^_di_, i.e., b_i_ = 1 and low *S*^2^_di_ scores are indicative of highly stable genotypes (Table 6). Genotypes G6, G8, G11, and G13 with b_i_ values > 1 and yield performance greater than the overall mean were adapted to the favorable environments. Genotypes with bi values < 1 and mean yields lower than the overall mean have poor adaptation and may have specific adaptation for low-yielding environments. Genotypes G11, G6, and G12, with mean grain yield ranks of 1, 2, and 3, and S^2^_di_ ranks of 3, 1, and 4, respectively, had a good combination of yield and stability statistics (Table 7). Based on the W^2^_i_ and σ^2^_i_, genotypes G6, G12, G11, and G2 had the lowest values and were identified as the most stable. The CV statistics identified genotypes G2, G7, G11, and G12 as the four best-ranked genotypes. Using ASV, the best-ranked genotypes with grain yield mean performance G11, G6, and G12 had higher ASV values and were ranked 9, 7, and 8, respectively. The other AMMI based stability statistics: the average of the squared EV, SIPC, and Za identified genotype G6 as the most stable genotype, and a different rank order for other high-yielding genotypes such as G11, G12, G2, and G13. Similarly, genotype G6 was found to be the most stable by the WAASB stability score, followed by G2 and G13, whereas the lowest yielding genotypes, G9, G5, and G4, were identified as the most unstable by WAASB. The BLUP-based stability parameter that considers stability, adaptability, and mean performance (HMRPGV) found a similar ranking of genotypes as mean grain yield (Tables 6 and 7).

**Table 6.** The mean grain yield (GY) and stability statistics values for 13 faba bean genotypes across 15 environments

| Statistics | Genotypes |  |  |  |  |  |  |  |  |  |  |  |  |
| --- | --- | --- | --- | --- | --- | --- | --- | --- | --- | --- | --- | --- | --- |
|  | G1 | G2 | G3 | G4 | G5 | G6 | G7 | G8 | G9 | G10 | G11 | G12 | G13 |
| GY | 2.31 | 2.44 | 2.20 | 2.12 | 2.11 | 2.53 | 2.23 | 2.36 | 2.03 | 2.16 | 2.53 | 2.50 | 2.41 |
| $W^2_i$ | 0.65 | 0.62 | 1.01 | 1.10 | 1.99 | 0.58 | 0.69 | 0.96 | 1.98 | 0.76 | 0.61 | 0.58 | 0.65 |
| $\sigma^2_i$ | 0.05 | 0.05 | 0.08 | 0.09 | 0.16 | 0.04 | 0.05 | 0.07 | 0.16 | 0.06 | 0.05 | 0.04 | 0.05 |
| $S^2_{di}$ | 0.09 | 0.09 | 0.14 | 0.14 | 0.28 | 0.06 | 0.07 | 0.11 | 0.27 | 0.11 | 0.09 | 0.08 | 0.09 |
| $b_i$ | 1.00 | 1.00 | 0.91 | 0.88 | 0.96 | 1.14 | 0.85 | 1.15 | 1.11 | 0.95 | 1.03 | 0.98 | 1.03 |
| CV | 33.08 | 31.16 | 32.61 | 32.82 | 37.78 | 33.83 | 29.14 | 37.04 | 43.78 | 33.89 | 30.88 | 29.88 | 32.39 |
| ASV | 0.55 | 0.17 | 0.17 | 0.34 | 1.05 | 0.34 | 0.20 | 0.59 | 1.02 | 0.28 | 0.54 | 0.50 | 0.25 |
| EV | 0.04 | 0.05 | 0.14 | 0.09 | 0.14 | 0.03 | 0.05 | 0.09 | 0.14 | 0.09 | 0.04 | 0.04 | 0.04 |
| SIPC | 0.96 | 0.95 | 1.75 | 1.43 | 1.88 | 0.73 | 1.14 | 1.44 | 2.06 | 1.57 | 1.10 | 1.00 | 0.95 |
| Za | 0.164 | 0.143 | 0.207 | 0.228 | 0.309 | 0.104 | 0.150 | 0.199 | 0.340 | 0.203 | 0.175 | 0.157 | 0.150 |
| WAASB | 0.17 | 0.16 | 0.21 | 0.24 | 0.28 | 0.15 | 0.17 | 0.23 | 0.33 | 0.21 | 0.19 | 0.18 | 0.17 |
| HMRPGV | 1.00 | 1.06 | 0.96 | 0.92 | 0.91 | 1.09 | 0.98 | 1.01 | 0.85 | 0.94 | 1.10 | 1.09 | 1.04 |
| $S^{(1)}$ | 3.90 | 3.43 | 4.38 | 3.60 | 3.43 | 2.32 | 3.71 | 4.13 | 4.29 | 3.26 | 2.36 | 2.67 | 4.11 |
| $S^{(2)}$ | 11.07 | 9.00 | 14.43 | 11.70 | 8.74 | 3.98 | 12.12 | 12.70 | 6.07 | 8.83 | 4.27 | 5.55 | 12.21 |
| $S^{(3)}$ | 23.01 | 14.00 | 33.67 | 36.12 | 23.54 | 5.65 | 27.67 | 27.48 | 55.31 | 26.87 | 5.71 | 7.88 | 21.55 |
| $S^{(6)}$ | 5.98 | 3.78 | 7.00 | 8.09 | 6.77 | 2.62 | 6.67 | 7.03 | 13.08 | 7.39 | 2.45 | 2.70 | 5.43 |
| NP<sup>(1)</sup> | 3.13 | 2.13 | 3.53 | 3.07 | 3.33 | 3.00 | 2.87 | 3.33 | 4.73 | 3.33 | 2.73 | 2.87 | 2.87 |
| NP<sup>(2)</sup> | 0.41 | 0.25 | 0.78 | 0.95 | 0.82 | 0.29 | 0.63 | 0.42 | 2.73 | 1.20 | 0.33 | 0.24 | 0.36 |
| NP<sup>(3)</sup> | 0.54 | 0.28 | 0.66 | 0.78 | 0.81 | 0.37 | 0.57 | 0.63 | 1.20 | 0.81 | 0.31 | 0.35 | 0.41 |
| NP<sup>(4)</sup> | 0.58 | 0.38 | 0.73 | 0.79 | 0.66 | 0.24 | 0.61 | 0.64 | 1.05 | 0.71 | 0.23 | 0.27 | 0.52 |
| KR | 12 | 8 | 19 | 22 | 25 | 3 | 15 | 15 | 25 | 18 | 4 | 5 | 11 |

See Tables 1 and 3 for genotypes and statistics names

**Table 7.** Rank of mean grain yield (GY), stability statistics and average sum of ranks (ASR) of all stability statistics for 13 faba bean genotypes tested from 15 environments

| Statistics | Genotypes |  |  |  |  |  |  |  |  |  |  |  |  |
| --- | --- | --- | --- | --- | --- | --- | --- | --- | --- | --- | --- | --- | --- |
|  | G1 | G2 | G3 | G4 | G5 | G6 | G7 | G8 | G9 | G10 | G11 | G12 | G13 |
| GY | 7 | 4 | 9 | 11 | 12 | 2 | 8 | 6 | 13 | 10 | 1 | 3 | 5 |
| $W^2_i$ | 5 | 4 | 10 | 11 | 13 | 1 | 7 | 9 | 12 | 8 | 3 | 2 | 6 |
| $\sigma^2_i$ | 5 | 4 | 10 | 11 | 13 | 1 | 7 | 9 | 12 | 8 | 3 | 2 | 6 |
| $b_i$ | 8 | 7 | 3 | 2 | 5 | 12 | 1 | 13 | 11 | 4 | 10 | 6 | 9 |
| $S^2_{di}$ | 7 | 5 | 10 | 11 | 13 | 1 | 2 | 9 | 12 | 8 | 4 | 3 | 6 |
| CV | 8 | 4 | 6 | 7 | 12 | 9 | 1 | 11 | 13 | 10 | 3 | 2 | 5 |
| ASV | 10 | 2 | 1 | 6 | 13 | 7 | 3 | 11 | 12 | 5 | 9 | 8 | 4 |
| EV | 2 | 6 | 12 | 8 | 11 | 1 | 7 | 9 | 13 | 10 | 4 | 3 | 5 |
| SIPC | 4 | 3 | 11 | 8 | 12 | 1 | 7 | 9 | 13 | 10 | 6 | 5 | 2 |
| Za | 6 | 2 | 10 | 11 | 12 | 1 | 4 | 8 | 13 | 9 | 7 | 5 | 3 |
| WAASB | 5 | 2 | 9 | 11 | 12 | 1 | 4 | 10 | 13 | 8 | 7 | 6 | 3 |
| HMRPGV | 7 | 4 | 9 | 11 | 12 | 2 | 8 | 6 | 13 | 10 | 1 | 3 | 5 |
| $S^{(1)}$ | 9 | 5 | 13 | 7 | 5 | 1 | 8 | 11 | 12 | 4 | 2 | 3 | 10 |
| $S^{(2)}$ | 7 | 6 | 12 | 8 | 4 | 1 | 9 | 11 | 13 | 5 | 2 | 3 | 10 |
| $S^{(3)}$ | 6 | 4 | 11 | 12 | 7 | 1 | 10 | 9 | 13 | 8 | 2 | 3 | 5 |
| $S^{(6)}$ | 6 | 4 | 9 | 12 | 8 | 2 | 7 | 10 | 13 | 11 | 1 | 3 | 5 |
| $NP^{(1)}$ | 8 | 1 | 12 | 7 | 9 | 6 | 3 | 9 | 13 | 9 | 2 | 3 | 3 |
| $NP^{(2)}$ | 6 | 2 | 9 | 11 | 10 | 3 | 8 | 7 | 13 | 12 | 4 | 1 | 5 |
| $NP^{(3)}$ | 6 | 1 | 9 | 10 | 12 | 4 | 7 | 8 | 13 | 11 | 2 | 3 | 5 |
| $NP^{(4)}$ | 6 | 4 | 11 | 12 | 9 | 2 | 7 | 8 | 13 | 10 | 1 | 3 | 5 |
| KR | 6 | 4 | 10 | 11 | 12 | 1 | 7 | 7 | 12 | 9 | 2 | 3 | 5 |
| ASR | 6.35 | 3.7 | 9.35 | 9.35 | 10.2 | 2.9 | 5.85 | 9.2 | 12.6 | 8.45 | 3.75 | 3.5 | 5.35 |

#### 3.6.2. Non-parametric measures of stability

According to stability statistics S^(1)^, S^(2)^, S^(3)^, and S^(6)^, genotypes G6, G11, and G12 had the lowest value in rank and are deemed as the most stable genotypes, while genotypes G3, G8, and G9 had relatively higher values of these statistics, indicating lower stability. The NP^(1)^, NP^(2)^, NP^(3)^, and NP^(4)^ considered genotypes G2, G6, G11, and G12 as more stable with a slight rank difference in the range of 1 to 4. However, the NP^(1)^ placed genotype G6 on a rank of six. Similarly, the KR stability index recognized genotypes G2, G6, G11, and G12 as the most stable. Overall, the results of non-parametric statistics were comparable to each other and identified genotypes G2, G6, G11, and G12 as stable genotypes.

### 3.7 Association among stability statistics

A heatmap of the Spearman’s rank correlation coefficient between mean grain yield (GY) and estimated stability parameters is displayed in Figure 2A. The results showed that GY was strongly and positively correlated with all other parametric and non-parametric indices, with the exception of b_i_, ASV, S^(1)^, and S^(2)^. Nevertheless, none of the estimated stability parameters were significantly associated with bi and ASV, except for ASV with CV. The non-parametric stability measures S^(1)^, and S^(2^ were only correlated with S^(3)^, S^(6)^, NP^(4)^, KR, W^2^_i_, σ^2^_i_, and EV. The CV indicated an association with all of the other stability measures evaluated except S^(1)^, S^(2)^, S^(3)^, b_i_, EV, and SIPC. However, the remaining stability parameters displayed a strong positive correlation with each other.

**Figure 2.**
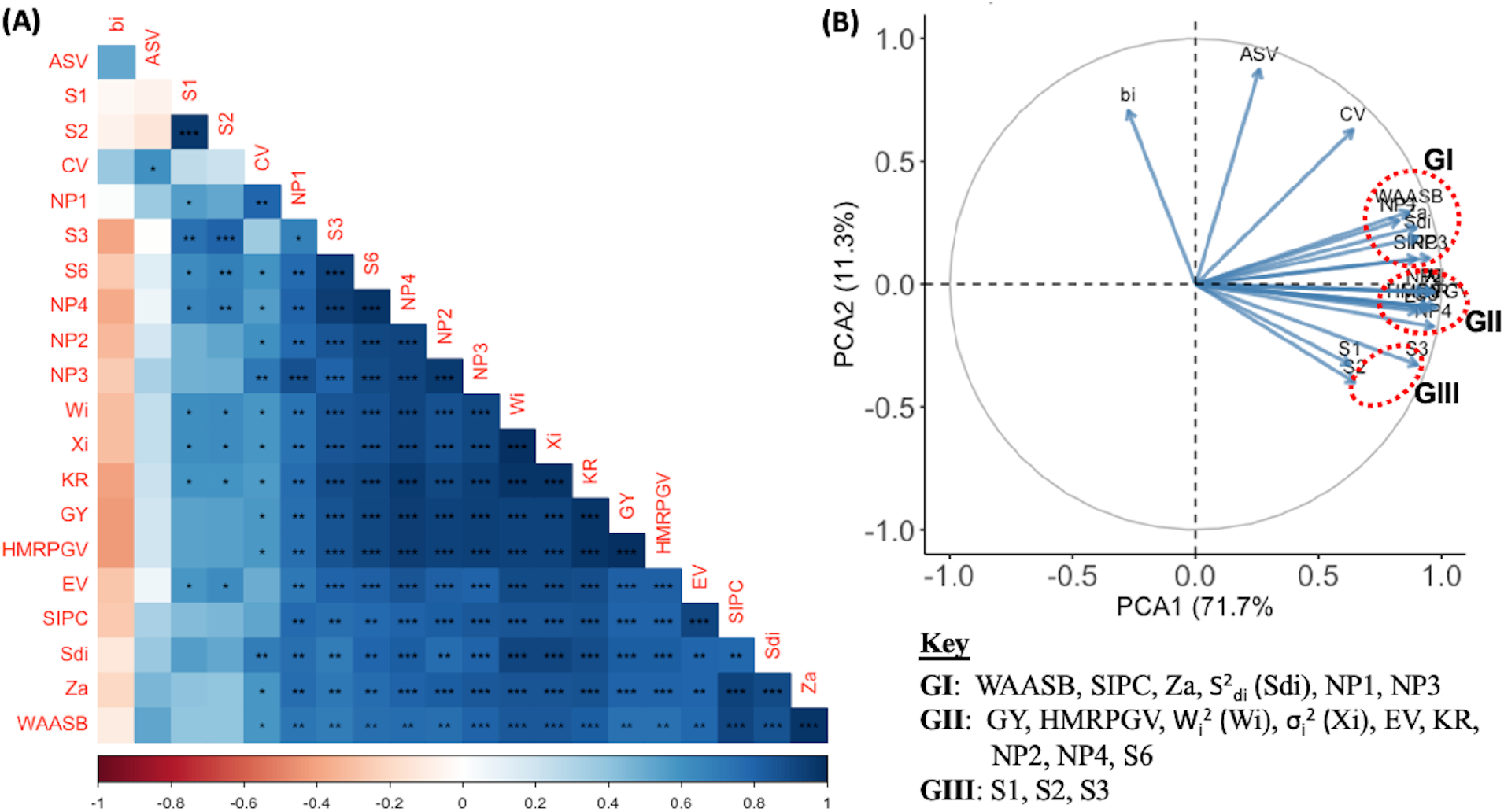
Heatmap of Spearman’s rank correlation (A) and biplot of principal component analysis (B) of mean yield (GY) and 20 stability statistics. *, **, and *** in the heatmap indicate significant at 0.05, 0.01, and 0.001 probability levels, respectively

The PCA based on the rank correlation matrix was performed to gain a better understanding of the interrelationships among the stability parameters. As shown in Figure 2B, the first two axes (PC1 and PC2) explained 82.7% of the total variance. The vectors of all of the indices were close to the edge of the circle, except b_i_, S^(1)^, and S^(2)^, indicating that they were well explained by the plane of factors. The stability parameters were graphically classified into distinct groups, with the cosine of the angle between their vectors approximating the association between each pair. The stability indices b_i_, ASV, and CV are placed separately from other stability measures, and each of them stands alone. The other remaining stability parameters were divided into three sub-groups (GI, GII, and GIII). Group I included the WAASB, SIPC, Za, S^2^_di_, NP^(1)^ and NP^(3)^, and the second group (II) contained GY, HMRPGV, W^2^_i_, σ^2^_i_, EV, KR, NP^(2)^, NP^(4)^, and S^(6)^, whereas the stability indices S^(1)^, S^(2)^, and S^(3)^ were classified in group III.

### 3.8 Clustering and ranking of genotypes

Hierarchical cluster analysis based on average sum rank (ASR) of stability measures and mean grain yield was computed to classify genotypes with similar performance regarding stability and productivity. The analysis grouped the 13 faba bean genotypes into two main clusters (Figure 3B). The first cluster was further subdivided into two subclusters, including genotypes G3, G10, G5, G4, and G9 in the first subcluster. This subgroup had a lower average grain yield (2.121 vs. 2.301 t ha^-1^) and the highest ASR values (Table 7). The second subcluster comprised three genotypes, G7, G1, and G8, and had an average grain yield equal to the overall mean (2.301 t ha^-1^) and higher ASR values for stability parameters. The other main cluster contains G2, G12, G13, G16, and G11, which had a higher average grain yield than the overall mean (2.480 vs. 2.301 t ha^-1^) and the lowest ASR values for stability parameters. This result was compared with the WAASB superiority index, WAASBY. The WAASBY values were calculated by considering the weights of 65 for grain yield and 35 for stability (WAASB). The genotypes with the highest WAASBY scores were G6 (99.11), followed by G11 (89.32), G12 (88.30), G2 (87.84) and G13 (81.16) (Figure 3A; Supplemental Table S4). The genotype with the lowest WAASBY score was G9 (0), followed by G5 (19.43), G4 (32.11), G10 (46.73), and G3 (50.14). These genotypes had the highest ASR values and the lowest average mean grain yield.

**Figure 3.**
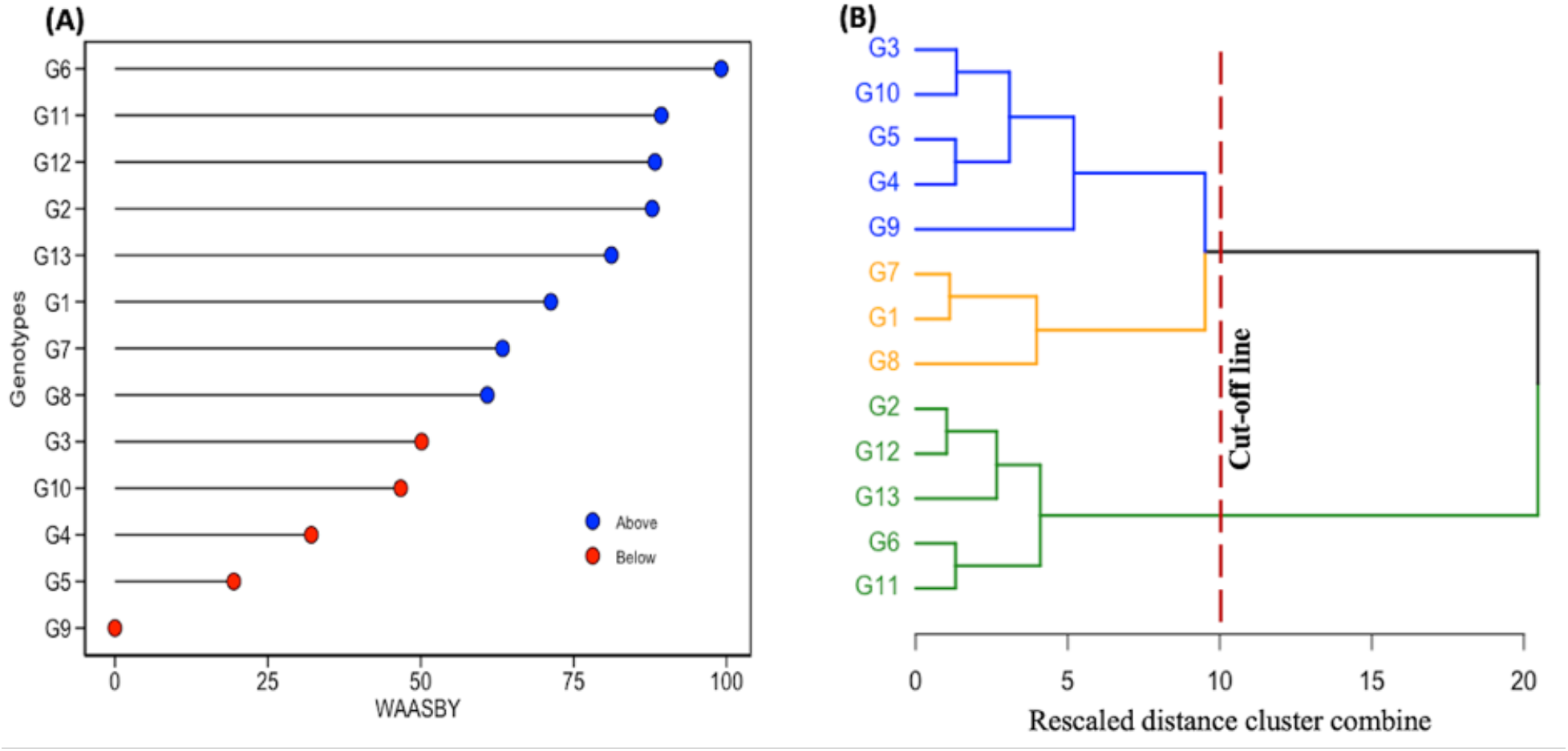
Estimated values of the weighted average of the stability (WAASB) and mean grain yield (WAASBY) (A), and a dendrogram showing the hierarchical classification of 13 evaluated faba bean genotypes based on ranks of mean grain yield and the average sum of ranks of all stability statistics (B). The WAASBY was computed considering the weights of 65 and 35 for yielding and stability, respectively. See Table 1 for genotype names.

## 4. DISCUSSION

World faba bean production has decreased by 56% over the past 50 years (FAOSTAT, 2021). In some aspects, the decline in production might be related to interannual yield instability (Cernay et al., 2015), the genotypes’ poor environmental adaptation (Zong et al., 2019), susceptibility to biotic stresses (Rubiales and Khazaei, 2022), the extensive practice of cereal monoculture in many countries and the use of chemical fertilizers (Jensen et al., 2010), and, of course, the crop’s antinutritional factors (Khazaei et al., 2019). So far, progressive efforts have been made to improve the nutritional quality; these traits have been incorporated into elite breeding lines, and the release of improved varieties has recently started in western Canada. Thus, it is essential to understand the magnitude of the GEI, one of the contributing factors to yield instability, before recommending new varieties for cultivation. In the present study, we employed multiple statistical models to investigate the grain yield performance and stability of faba bean genotypes.

The current study found highly significant differences between genotypes, environments, and GEI effects. Our results revealed that environmental changes had the largest effect on grain yield, resulting in lower heritability. The result is consistent with previous research showing that the environment has a significant impact on faba bean grain production (Fikere et al., 2008; Flores et al., 2012; Temesgen et al., 2015; Skovbjerg et al., 2020; Papastylianou et al., 2021). As shown in Supplemental Figure 1A, the productivity of the environment was highly variable, ranging from 0.881 to 3.514 t ha^-1^, with a difference greater than the grand mean. The predominant environmental effects on faba bean grain yield are attributed to the prevalence of abiotic and biotic factors due to climatic and edaphic variation (Table 2). These factors, especially if they occur during the reproductive stage of the crop, i.e., flowering, podding, and grain filling, can significantly reduce grain yield (e.g., Link et al., 1999; Mwanamwenge et al., 1999; Katerji et al., 2011; Ammar et al., 2015). Significant GEI effects that are greater than those contributed by genotype also imply that genotype responses differ across test environments, which in turn suggests a significant difference in genotypic performances and their rank orders. Likewise, Link et al. (1996), Annicchiarico and Iannucci (2008), and Papastylianou et al. (2021) reported a large crossover GEI in faba bean METs. This phenomenon could reduce the accuracy of selection for grain yield and obstruct the development of new cultivars due to the masking effects of variable environments (Kang and Pham, 1991). Therefore, accurate prediction models must be used to correctly analyze and interpret the yield performance, adaptability patterns, and stability of genotypes in METs (Gauch and Zobel, 1988).

The AMMI is the most frequently used model in the partitioning of GEI into IPCs in METs. The AMMI analysis of this study showed that the first five IPCs are significant, and the first two IPCs accounted for over half of the total GEI. The interpretation of the AMMI analysis based on the first two components could be biased as only half of the variation is exploited. In this situation, employing model diagnosis holds the highest importance for choosing the best model for the data set (Gauch, 2013). Importantly, the AMMI is not just one model; it is rather a series of models, ranging from AMMI0 to AMMIF. Consequently, we evaluated the AMMI family and BLUP models, and the BLUP methods were found to outperform the AMMI models in predicting genotypic response. Our findings are consistent with the results of Piepho (1994), who demonstrated that the BLUP performs better than any member of the AMMI family in predicting faba bean yield in MET. Similar results are also reported for other crops (van Eeuwijk et al., 2016; Olivoto et al., 2019; Huang et al., 2021; Nataraj et al., 2021). Nevertheless, there are many cases where the AMMI model has been used even though the proportion of GEI explained by the first two IPCs was low (Tigabu et al., 2017; Bocianowski et al., 2019; Pour-Aboughadareh et al., 2022b). In the current study, to leverage the advantages of both models, we used the WAASB, which incorporates all IPCs from the AMMI into BLUP methods to properly quantify the genotypic stability. The WAASB biplot is a useful tool for simultaneously examining yield performance and stability as it provides important details regarding the distribution of genotypes and environments. The results revealed genotypes G6 (AO1155), G11 (1089-1-2), G12 (1310-5), G13 (1239-1), and G2 (DL Tesoro) as stable and high-yielding genotypes. Furthermore, the environments E5 (Roblin 2020), E7 (Stonewall 2020), and E15 (Melfort 2019) were identified as having good genotype discrimination ability. Many studies used the WAASB to identify genotypes that are highly productive and widely adaptable in various crops (Huang et al., 2021; Nataraj et al., 2021; Koundinya et al., 2021; Pour-Aboughadareh et al., 2022b).

Numerous stability statistics and models have been presented for assessing the stability of the tested genotypes in METs. In this study, we used several parametric and non-parametric stability statistics to better understand the stability of faba bean genotypes. Stability methods are commonly classified as static or dynamic concepts, depending on their relationship with yield performance (Leon, 1985). The static stability or biological concepts state that a stable genotype maintains a constant yield regardless of diverse environments and its yield performance has an environmental variance near to zero (Becker and Leon, 1988). The dynamic stability, or agronomic concepts, implies that a genotype’s performance responds consistently to environmental changes with the same trend as the mean response of the tested genotypes, i.e., no GEI. In contrast to the static stability measure, the dynamic stability measure is dependent on a specific set of genotypes that have been evaluated (Lin et al., 1986). Moreover, the classification of stability parameters into static and dynamic concepts depends on the nature of the data and test environments that determine their association with yield performance (Pour-Aboughadareh et al., 2022a). In this regard, we performed Spearman’s rank correlation and PCA (see Figure 2 A & B) for further dissection of the relationships among stability statistics and the stability concepts.

Our findings revealed that the GY was strongly and positively correlated with all stability statistics, with the exception of b_i_, ASV, S^(1)^, and S^(2)^. None of the estimated stability parameters were significantly associated with bi and ASV, except for ASV with CV. However, the majority of the stability parameters calculated in this study displayed a strong positive correlation with each other. Our results also showed that the PCA biplot depicts the b_i_, ASV, and CV separated from the other groups, and each stand alone, representing the measure of stability in a static sense. These statistics were not significantly correlated with mean grain yield, except CV, and they might be applied to identify genotypes adapted to environments with unfavourable growing conditions. Group I included WAASB, SIPC, Za, S^2^_di_, NP^(1)^, and NP^(3)^ that were influenced simultaneously by both grain yield and stability. It was found that genotypes identified using these methods had average stability. However, these genotypes could not perform as good as the responsive ones in a favourable environment. Group II consisted of the GY with HMRPGV, NP^(2)^, NP^(4)^, σ^2^_i_, W^2^_i_, KR, EV, and S^(6)^. These statistics correspond to the dynamic concept of stability and favour selection based on grain yield (Becker and Leon, 1988), and can be used to identify genotypes that are adapted to favorable conditions. The high-yielding genotypes G11, G6, G12, and G2 had a rank in the range of 1 to 4 with these stability statistics, showing that the grain yield had a main influence on the rankings of genotypes. Our results also showed that S^(1)^, S^(2)^, and S^(3)^ were included in group III. Like group I stability parameters, they are related to the static concept. All the stability methods included in groups I and II had a strong positive association with each other and mean grain yield, although there is inconsistency in ranking patterns in the selection of stable genotypes.

Our findings also showed that some lines exhibited remarkable stable yield performance for some stability parameters and instability for others. This is one of the problems that has been identified in GE interaction studies (Khalili and Pour-Aboughadareh, 2016). This problem could be solved using the ASR values of the calculated stability statistics (Alizadeh et al., 2022; Pour-Aboughadareh et al., 2022b). The low ASR value indicates a high level of stability; therefore, genotypes G6, G12, G2, G11, and G13 are identified as the most stable genotypes in this study. Furthermore, HCA based on the ASR values and mean grain yield was used to cluster the genotypes into qualitatively homogeneous high-yielding and stable subsets (Lin et al., 1986; Becker and Leon, 1988). Accordingly, the 13 faba bean test genotypes were divided into two main clusters. The first cluster was further subdivided into two subclusters, with the first subcluster consisting of genotypes that had a mean grain yield lower than the overall mean and the highest ASR values. The other subcluster included genotypes that had a mean grain yield above the overall mean as well as relatively higher ASR values for stability parameters. Some of the genotypes in this subcluster may have specific adaptations to some of the environments, as shown in Figure 2. The second main cluster comprised high-yielding genotypes with a low ASR value of stability parameters (ranked from 1 to 5), identified as high-yielding and more stable genotypes.

Finally, we compared the results of ASR values of the stability parameters with WAASBY in identifying high-yielding stable genotypes of faba bean. To determine the efficiency and suitability of the WAASB statistics in identifying the ideal faba bean genotypes, as previously stated in the objectives. Like the ASR values, the WAASBY index identified genotype G6 as the most high-yielding and stable, followed by G11, G12, G2, and G13. The superiority index, WAASBY, may be more advantageous as it allows weighting between performance of response variables and the WAASB stability score for simultaneous selection of stability and productivity under a mixed effect model (Olivoto et al., 2019). Therefore, breeders can prioritize weights for mean grain yield and stability as per their breeding objectives and cultivar recommendations. High stability is only advantageous when associated with high yielding performance, and it is the least desirable when combined with low performance (Yan et al., 2007). In the current study, the WAASBY index was computed by assigning weights of 65 and 35, respectively, for grain yield and stability. Hence, the genotype G6 was found to have the highest superiority index, with a grain yield greater than the overall mean, and can be used for the improvement of adaptation in faba bean breeding programs. In Europe, faba bean synthetic lines have shown to have better yield stability than lines developed by recurrent/mass selection or pedigree selection (e.g., Stelling et al., 1994; Skovbjerg et al., 2020). However, in our study, this was not the case. The main reason is that most synthetic lines used in this study were bred by NPZ (Norddeutsche Pflanzenzücht, Germany) and may have less adaptability to the western Canada climate.

## 5 CONCLUSIONS

In the current study, 13 faba bean genotypes were tested across 15 environments in western Canada to exploit the effects of GE interaction and simultaneous selection of the best genotypes for mean grain yield and stability. The AMMI model and BLUP method demonstrated that the grain yield was highly affected by the genotype, environment, and their interaction. The combination of the AMMI and BLUP methods made it possible to dissect GEI effects more accurately and the suitability of the WAASB in multi-environment experiments in faba bean. Fifteen of the 20 stability statistics revealed a significant positive correlation with grain yield, and most of the statistics were found to be positively and significantly correlated with each other. This result indicated that non-parametric statistics seem to be useful alternatives to complement parametric methods for identifying the most stable genotypes. Both univariate and multivariate statistical groups identified genotypes G6 (AO1155), G11 (1089-1-2) and G12 (1310-5) as more high-yielding and stable genotypes than the best check G2 (DL Tesoro). This result is confirmed with the WAASBY index, indicating the efficiency of the WAASBY statistics in selecting superior faba bean genotypes. Overall, the genotype G6 (AO1155) with the highest yielding and stable performance could be the most promising genetic resource for improving and stabilising faba bean grain yield in western Canada.

## ACKNOWLEDGMENTS

The financial support of the Saskatchewan Agriculture Development Fund, Saskatchewan Pulse Crop Development Board and NSERC is greatly appreciated. We thank Lana Shaw (South East Research Farm), Jessica Frey (PCDF/Manitoba Agriculture), Jesse Mutcheson (DL Seeds), Christy Hoy (Alberta Agriculture), and Kevin Baron (Solum Valley Biosciences) for their kind assistance with field trials.

## AUTHOR CONTRIBUTIONS

Tadese S. Gela: Conceptualization; Data curation; Formal analysis; Validation; Methodology; Writing-original draft. Hamid Khazaei: Conceptualization; Data curation; Writing-review & editing. Rajib Podder: Methodology. Albert Vandenberg: Conceptualization; Resources; Writing-review & editing.

## Supplemental materials

**Supplemental Table S1.** Analysis of variance of grain yield for individual environments.

| Environment | Mean Square |  |  | Mean grain<br>yield (t/ha) | CV(%) | h<sup>2</sup> | AS |
| --- | --- | --- | --- | --- | --- | --- | --- |
|  | Block | Genotype | Error |  |  |  |  |
| E1 | 0.02<sup>ns</sup> | 0.07* | 0.02 | 1.62 | 9.66 | 0.65 | 0.80 |
| E2 | 1.11** | 0.47* | 0.16 | 2.59 | 15.21 | 0.67 | 0.82 |
| E3 | 0.04<sup>ns</sup> | 0.12*** | 0.01 | 1.48 | 8.10 | 0.88 | 0.94 |
| E4 | 0.07<sup>ns</sup> | 0.62*** | 0.07 | 2.48 | 10.50 | 0.89 | 0.94 |
| E5 | 0.21<sup>ns</sup> | 0.22<sup>ns</sup> | 0.17 | 3.04 | 13.40 | 0.26 | 0.51 |
| E6 | 1.08** | 0.48** | 0.12 | 2.29 | 15.31 | 0.74 | 0.86 |
| E7 | 0.17** | 0.39*** | 0.03 | 2.98 | 5.46 | 0.93 | 0.97 |
| E8 | 0.02<sup>ns</sup> | 0.11*** | 0.01 | 1.56 | 7.63 | 0.87 | 0.93 |
| E9 | 0.38*** | 0.25*** | 0.03 | 1.80 | 8.85 | 0.90 | 0.95 |
| E10 | 0.01<sup>ns</sup> | 0.05** | 0.01 | 0.88 | 13.37 | 0.75 | 0.86 |
| E11 | 0.62* | 0.43** | 0.13 | 2.93 | 12.39 | 0.70 | 0.83 |
| E12 | 0.19<sup>ns</sup> | 0.75** | 0.16 | 2.79 | 14.24 | 0.79 | 0.89 |
| E13 | 0.53<sup>ns</sup> | 0.21<sup>ns</sup> | 0.16 | 3.51 | 11.25 | 0.27 | 0.52 |
| E14 | 0.01<sup>ns</sup> | 0.12** | 0.03 | 1.91 | 9.18 | 0.74 | 0.86 |
| E15 | 0.25** | 0.09* | 0.04 | 2.64 | 7.23 | 0.60 | 0.77 |
| DF | 2 | 12 | 24 |  |  |  |  |

\*, \*\* and \*\*\*, significant at the 0.05, 0.01 and 0.001 probability levels, respectively; ns, non-significant, CV, coefficient of variation; h<sup>2</sup>, broad-sense heritability; AS, accuracy of selection; DF, degrees of freedom.

**Supplemental Table S2.**
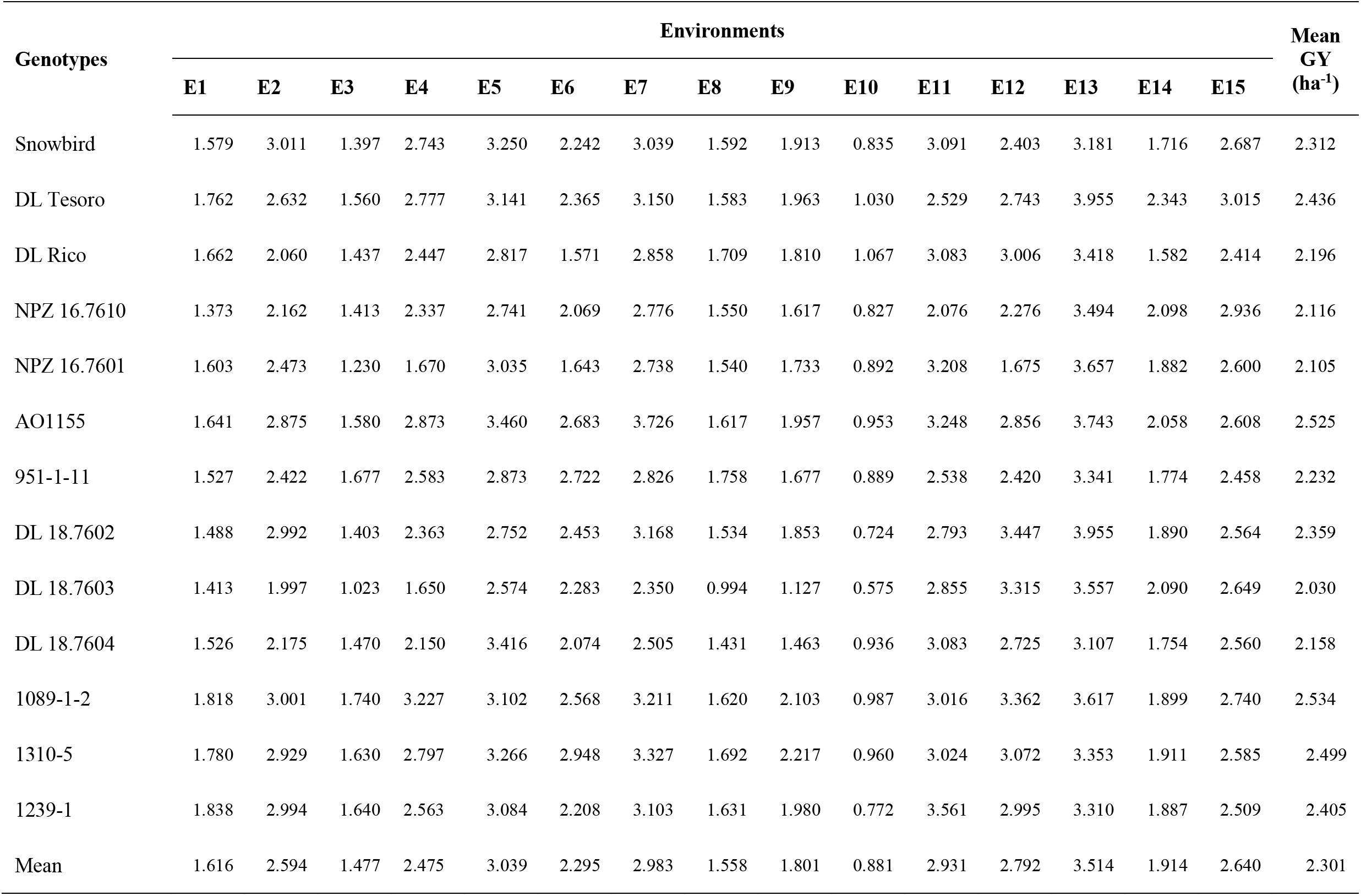
Mean grain yield (GY; t ha^−1^) of 13 faba bean genotypes tested in 15 environments during the 2019–2020 cropping season

| Genotypes | Environments |  |  |  |  |  |  |  |  |  |  |  |  |  |  | Mean<br>GY<br>(ha<sup>-1</sup>) |
| --- | --- | --- | --- | --- | --- | --- | --- | --- | --- | --- | --- | --- | --- | --- | --- | --- |
|  | E1 | E2 | E3 | E4 | E5 | E6 | E7 | E8 | E9 | E10 | E11 | E12 | E13 | E14 | E15 |  |
| Snowbird | 1.579 | 3.011 | 1.397 | 2.743 | 3.250 | 2.242 | 3.039 | 1.592 | 1.913 | 0.835 | 3.091 | 2.403 | 3.181 | 1.716 | 2.687 | 2.312 |
| DL Tesoro | 1.762 | 2.632 | 1.560 | 2.777 | 3.141 | 2.365 | 3.150 | 1.583 | 1.963 | 1.030 | 2.529 | 2.743 | 3.955 | 2.343 | 3.015 | 2.436 |
| DL Rico | 1.662 | 2.060 | 1.437 | 2.447 | 2.817 | 1.571 | 2.858 | 1.709 | 1.810 | 1.067 | 3.083 | 3.006 | 3.418 | 1.582 | 2.414 | 2.196 |
| NPZ 16.7610 | 1.373 | 2.162 | 1.413 | 2.337 | 2.741 | 2.069 | 2.776 | 1.550 | 1.617 | 0.827 | 2.076 | 2.276 | 3.494 | 2.098 | 2.936 | 2.116 |
| NPZ 16.7601 | 1.603 | 2.473 | 1.230 | 1.670 | 3.035 | 1.643 | 2.738 | 1.540 | 1.733 | 0.892 | 3.208 | 1.675 | 3.657 | 1.882 | 2.600 | 2.105 |
| AO1155 | 1.641 | 2.875 | 1.580 | 2.873 | 3.460 | 2.683 | 3.726 | 1.617 | 1.957 | 0.953 | 3.248 | 2.856 | 3.743 | 2.058 | 2.608 | 2.525 |
| 951-1-11 | 1.527 | 2.422 | 1.677 | 2.583 | 2.873 | 2.722 | 2.826 | 1.758 | 1.677 | 0.889 | 2.538 | 2.420 | 3.341 | 1.774 | 2.458 | 2.232 |
| DL 18.7602 | 1.488 | 2.992 | 1.403 | 2.363 | 2.752 | 2.453 | 3.168 | 1.534 | 1.853 | 0.724 | 2.793 | 3.447 | 3.955 | 1.890 | 2.564 | 2.359 |
| DL 18.7603 | 1.413 | 1.997 | 1.023 | 1.650 | 2.574 | 2.283 | 2.350 | 0.994 | 1.127 | 0.575 | 2.855 | 3.315 | 3.557 | 2.090 | 2.649 | 2.030 |
| DL 18.7604 | 1.526 | 2.175 | 1.470 | 2.150 | 3.416 | 2.074 | 2.505 | 1.431 | 1.463 | 0.936 | 3.083 | 2.725 | 3.107 | 1.754 | 2.560 | 2.158 |
| 1089-1-2 | 1.818 | 3.001 | 1.740 | 3.227 | 3.102 | 2.568 | 3.211 | 1.620 | 2.103 | 0.987 | 3.016 | 3.362 | 3.617 | 1.899 | 2.740 | 2.534 |
| 1310-5 | 1.780 | 2.929 | 1.630 | 2.797 | 3.266 | 2.948 | 3.327 | 1.692 | 2.217 | 0.960 | 3.024 | 3.072 | 3.353 | 1.911 | 2.585 | 2.499 |
| 1239-1 | 1.838 | 2.994 | 1.640 | 2.563 | 3.084 | 2.208 | 3.103 | 1.631 | 1.980 | 0.772 | 3.561 | 2.995 | 3.310 | 1.887 | 2.509 | 2.405 |
| Mean | 1.616 | 2.594 | 1.477 | 2.475 | 3.039 | 2.295 | 2.983 | 1.558 | 1.801 | 0.881 | 2.931 | 2.792 | 3.514 | 1.914 | 2.640 | 2.301 |

**Supplemental Table S3.** Predicted grain yield, best linear unbiased prediction (BLUP) values, and rank for 13 faba bean genotypes evaluated. The lower and upper limits represent the 95% confidence interval of prediction considering a two-tailed t-test

| Genotype | Genotype code | BLUP value | Predicted mean | Rank of genotype | Lower limit | Upper limit |
| --- | --- | --- | --- | --- | --- | --- |
| Snowbird | G1 | 0.009 | 2.310 | 7 | 2.164 | 2.457 |
| DL Tesoro | G2 | 0.114 | 2.415 | 4 | 2.268 | 2.561 |
| DL Rico | G3 | -0.088 | 2.213 | 9 | 2.067 | 2.359 |
| NPZ 16.7610 | G4 | -0.155 | 2.146 | 11 | 2.000 | 2.293 |
| NPZ 16.7601 | G5 | -0.164 | 2.137 | 12 | 1.991 | 2.283 |
| AO1155 | G6 | 0.188 | 2.489 | 2 | 2.342 | 2.635 |
| 951-1-11 | G7 | -0.057 | 2.243 | 8 | 2.097 | 2.390 |
| DL 18.7602 | G8 | 0.049 | 2.349 | 6 | 2.203 | 2.496 |
| DL 18.7603 | G9 | -0.227 | 2.074 | 13 | 1.927 | 2.220 |
| DL 18.7604 | G10 | -0.119 | 2.181 | 10 | 2.035 | 2.328 |
| 1089-1-2 | G11 | 0.196 | 2.496 | 1 | 2.350 | 2.643 |
| 1310-5 | G12 | 0.166 | 2.467 | 3 | 2.321 | 2.614 |
| 1239-1 | G13 | 0.087 | 2.388 | 5 | 2.242 | 2.535 |

**Supplemental Table S4.**
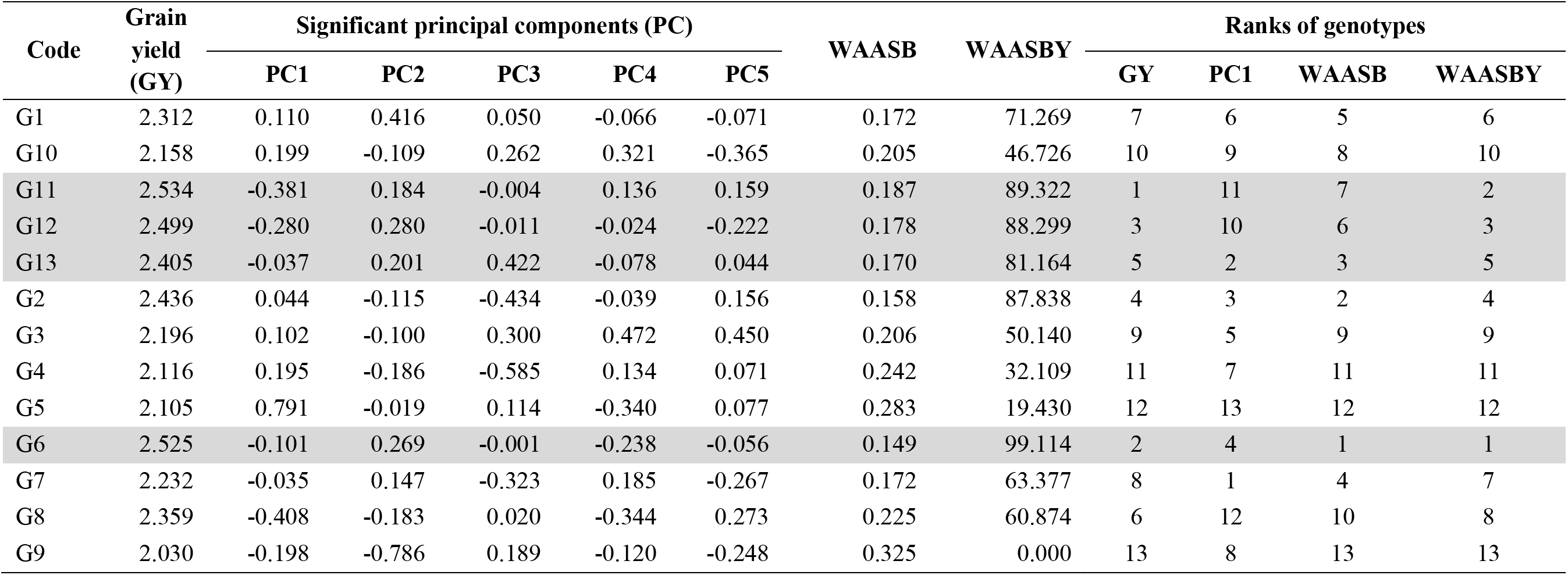
Results for WAASB estimation and ranks of the 13 faba bean genotypes evaluated in 15 environments

| Code | Grain yield (GY) | Significant principal components (PC) |  |  |  |  | WAASB | WAASBY | Ranks of genotypes |  |  |  |
| --- | --- | --- | --- | --- | --- | --- | --- | --- | --- | --- | --- | --- |
|  |  | PC1 | PC2 | PC3 | PC4 | PC5 |  |  | GY | PC1 | WAASB | WAASBY |
| G1 | 2.312 | 0.110 | 0.416 | 0.050 | -0.066 | -0.071 | 0.172 | 71.269 | 7 | 6 | 5 | 6 |
| G10 | 2.158 | 0.199 | -0.109 | 0.262 | 0.321 | -0.365 | 0.205 | 46.726 | 10 | 9 | 8 | 10 |
| G11 | 2.534 | -0.381 | 0.184 | -0.004 | 0.136 | 0.159 | 0.187 | 89.322 | 1 | 11 | 7 | 2 |
| G12 | 2.499 | -0.280 | 0.280 | -0.011 | -0.024 | -0.222 | 0.178 | 88.299 | 3 | 10 | 6 | 3 |
| G13 | 2.405 | -0.037 | 0.201 | 0.422 | -0.078 | 0.044 | 0.170 | 81.164 | 5 | 2 | 3 | 5 |
| G2 | 2.436 | 0.044 | -0.115 | -0.434 | -0.039 | 0.156 | 0.158 | 87.838 | 4 | 3 | 2 | 4 |
| G3 | 2.196 | 0.102 | -0.100 | 0.300 | 0.472 | 0.450 | 0.206 | 50.140 | 9 | 5 | 9 | 9 |
| G4 | 2.116 | 0.195 | -0.186 | -0.585 | 0.134 | 0.071 | 0.242 | 32.109 | 11 | 7 | 11 | 11 |
| G5 | 2.105 | 0.791 | -0.019 | 0.114 | -0.340 | 0.077 | 0.283 | 19.430 | 12 | 13 | 12 | 12 |
| G6 | 2.525 | -0.101 | 0.269 | -0.001 | -0.238 | -0.056 | 0.149 | 99.114 | 2 | 4 | 1 | 1 |
| G7 | 2.232 | -0.035 | 0.147 | -0.323 | 0.185 | -0.267 | 0.172 | 63.377 | 8 | 1 | 4 | 7 |
| G8 | 2.359 | -0.408 | -0.183 | 0.020 | -0.344 | 0.273 | 0.225 | 60.874 | 6 | 12 | 10 | 8 |
| G9 | 2.030 | -0.198 | -0.786 | 0.189 | -0.120 | -0.248 | 0.325 | 0.000 | 13 | 8 | 13 | 13 |

## Supplemental Figures

**Supplemental Figure S1.**
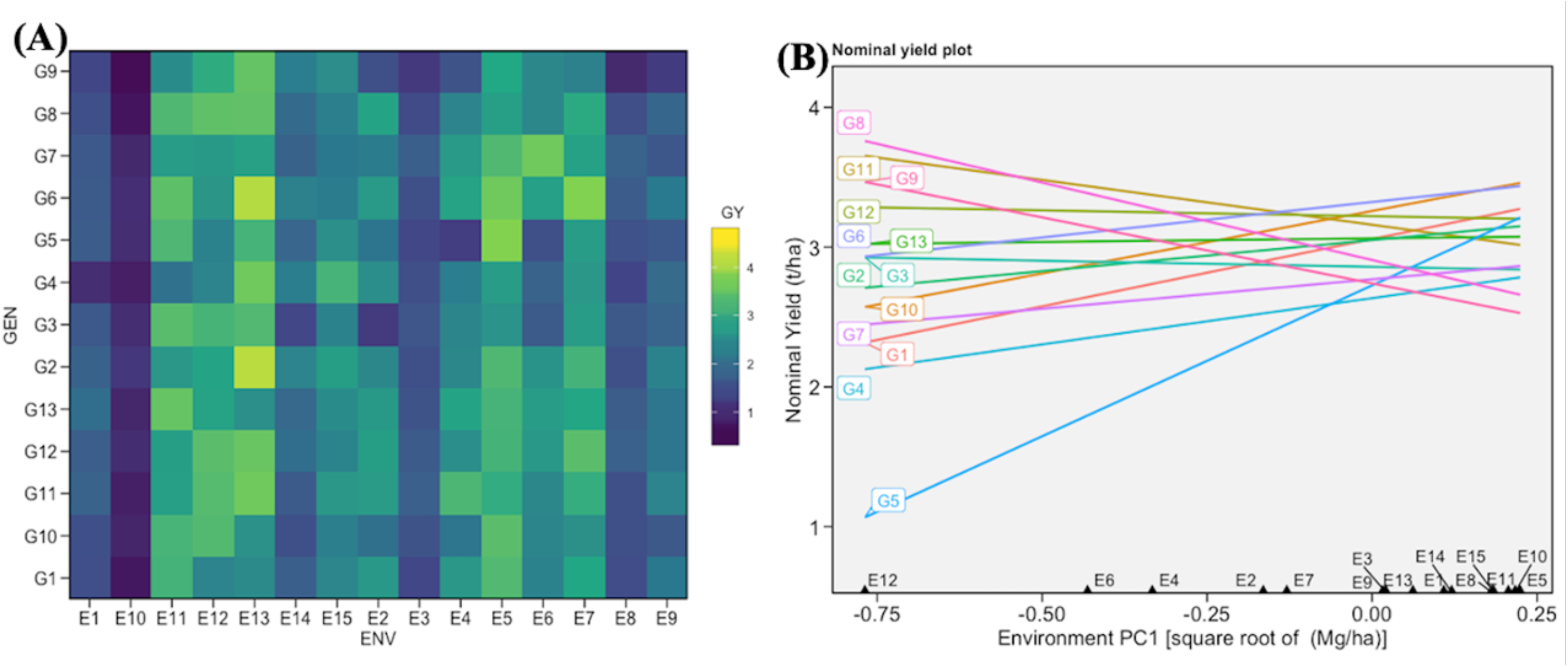
The mean grain yield (GY) variation of 13 faba bean genotypes across 15 environments (A) and a nominal grain yield describing the “which-won-where” view for the 13 faba bean genotypes as a function of the environment scores of the first interaction principal component axis (IPCA1) (B)

**Supplemental Figure S2.**
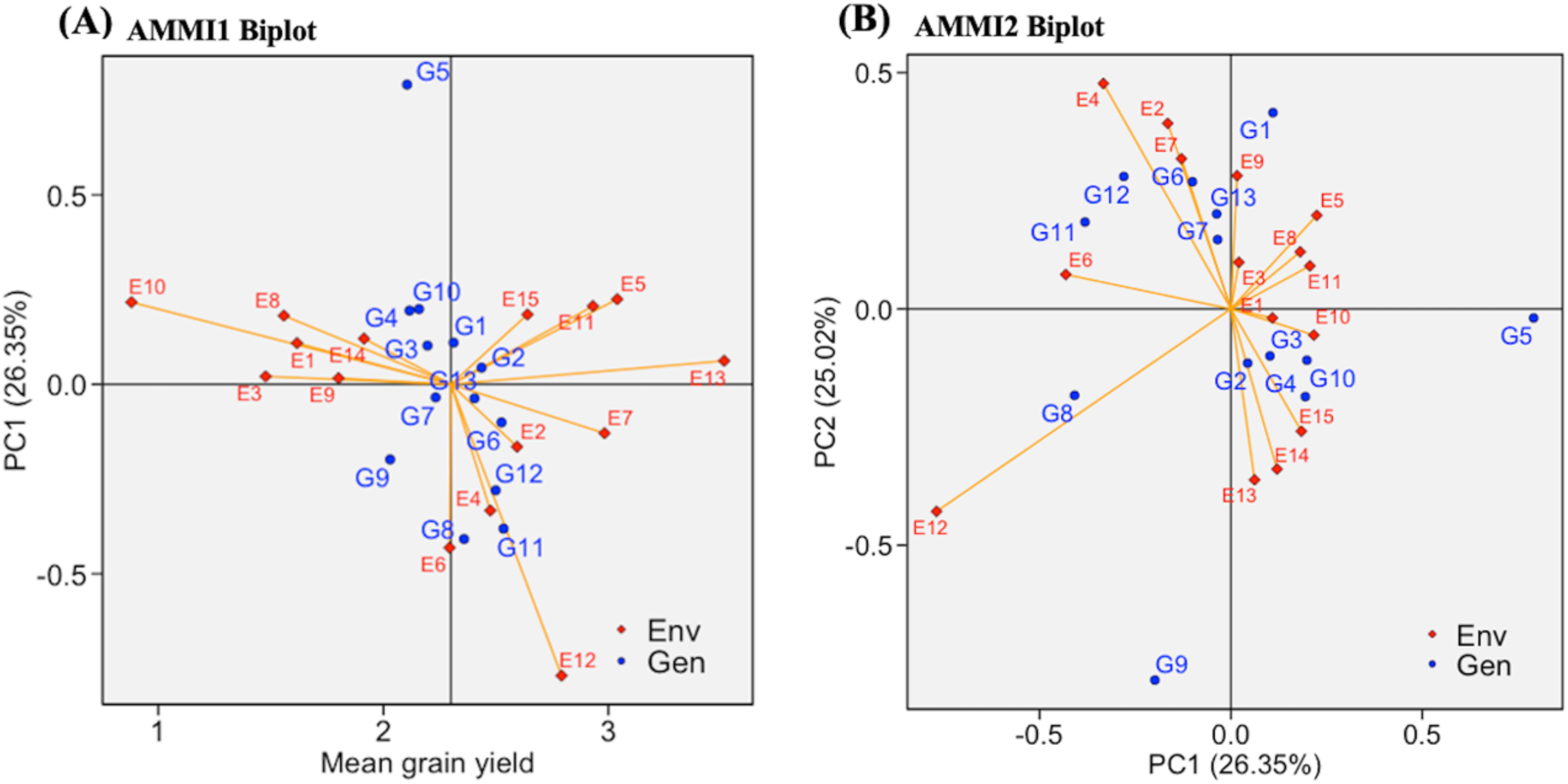
The AMMI1 (A) and AMMI2 (B) biplots indicate genotype by environment interaction for 13 faba bean genotypes evaluated in 15 environments. The genotype and environment codes are represented with blue and red icons, respectively. The Names of genotypes and environments are as defined in Table 1 and 2

**Supplemental Figure S3.**
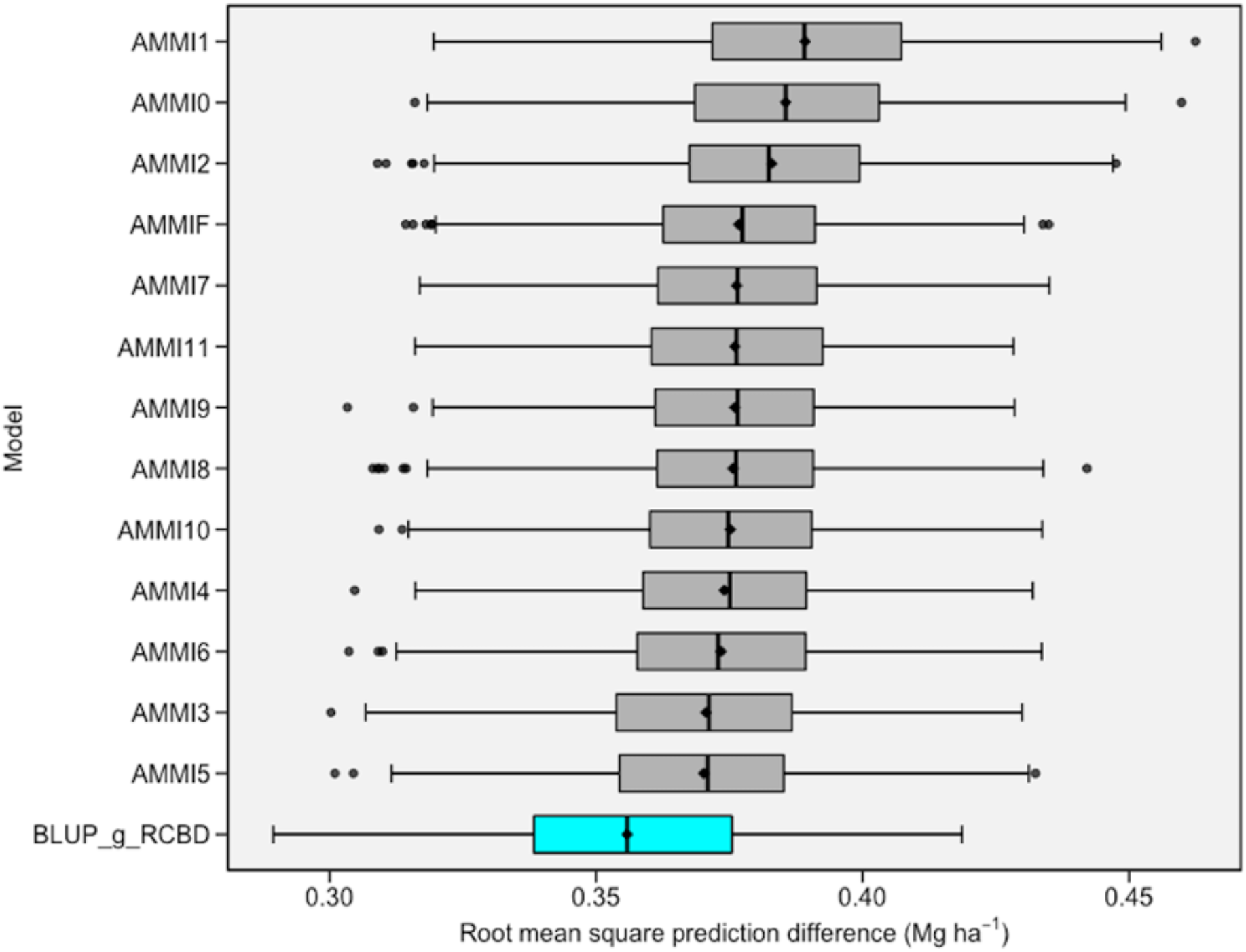
The boxplots display the distribution of the root mean square prediction difference (RMSPD) estimates used to assess the predictive accuracy of the additive main effects and multiplicative interaction (AMMI) family and best linear unbiased prediction (BLUP) for 13 faba bean genotypes evaluated in 15 environments. The RMSPD data was generated with 1000 bootstrapping

